# Optimizing chemo-scheduling based on tumor growth rates

**DOI:** 10.1101/263327

**Authors:** Jeffrey West, Paul Newton

**Affiliations:** Department of Aerospace & Mechanical Engineering, University of Southern California, CA 90089; Department of Aerospace & Mechanical Engineering, Mathematics, and Norris Comprehensive Cancer Center,University of Southern California, CA 90089

## Abstract

We review the classic tumor growth and regression laws of Skipper and Schable based on fixed exponential growth assumptions, and Norton and Simon’s law based on a Gompertzian growth assumption. We then discuss ways to optimize chemotherapeutic scheduling using a Moran process evolutionary game-theory model of tumor growth that incorporates more general dynamical and evolutionary features of tumor cell kinetics. Using this model, and employing the quantitative notion of Shannon entropy which assigns high values to low-dose metronomic (LDM) therapies, and low values to maximum tolerated dose (MTD) therapies, we show that low-dose metronomic strategies can outperform maximum tolerated dose strategies, particularly for faster growing tumors. The general concept of designing different chemotherapeutic strategies for tumors with different growth characteristics is discussed.

## I. INTRODUCTION TO TYPE OF PROBLEM IN CANCER

The most influential quantitative mathematical model of tumor growth and chemotherapeutic response was developed by Skipper and Schabel [1] in the 1960-70s. Their model was based on a series of careful *in vitro* experiments carried out using the L1210 mouse leukemia cells lines in which they observed that the cell numbers doubled at a fixed rate, implying constant exponential growth [2]. Thus, *Skipper’s first law* states that the doubling time of proliferating cancer cells is constant, forming a straight line on a semi-log plot [3]. A more subtle consequence of fixed exponential growth is encapsulated in *Skipper’s second law* which states that cell kill by chemotherapeutic agents follows first order kinetics. This implies that the percent of cells killed at a given drug dose is constant, proportional to the growth rate, regardless of tumor size or overall tumor burden [4]. If a given dose of drug reduces the number of cancer cells from 10^6^ to 10^5^ (killing 900,000 cells, or 90% reduction), the same therapy acting on 10^4^ cells will reduce it to 10^3^ cells (killing 9000 cells, or 90% reduction). Known as the *log-kill law* of chemotherapeutic response [5] (one log-kill is equivalent to 90% reduction), it has important therapeutic implications. If repeated, the third dose reduces the remaining 103 cells down to 102 cells (killing 900 cells), the fourth from 10^2^ to 10 (killing 90 cells), and the fifth from 10 to 1 (killing 9 cells). Thus, each round of identical chemotherapy kills fewer and fewer *numbers* of cells, making complete eradication of the tumor increasingly difficult, independent of any other effects. These two laws are all simple consequences of the hypothesis of constant exponential growth of the tumor volume (which typically becomes visible at roughly 1cm^3^, or 10^9^ tumor cells), 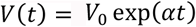, with fixed growth rate α. A third simple consequence is that tumors with higher growth rates (larger values of α) should also have steeper regression. This is evident in the regression curves of Figure 1 which have smaller negative slope higher up on the growth curve (slower growth regions) than lower on the growth curve.

**Figure 1:**
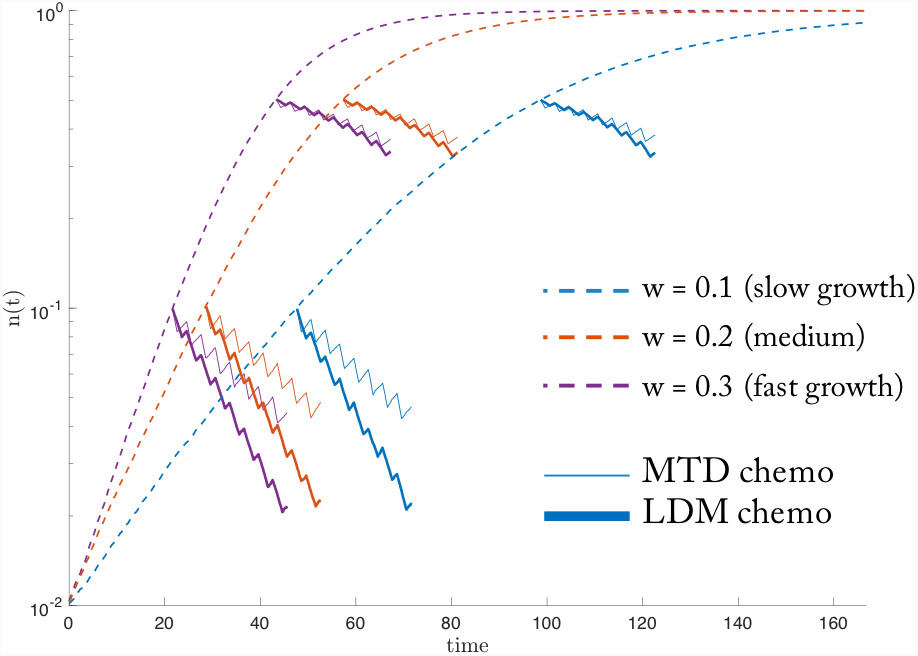
Tumor growth and regression. *Averages of 50 stochastic simulations of slow (w = 0.1; blue), medium (w = 0.2; red), and fast (w = 0.3; purple) growing tumors shown in dashed lines. Therapy is administered at identical tumor burden (solid lines) to compare 8 cycles of MTD (thin lines) and LDM (thick lines) chemotherapy schedules with equivalent total dose. For all growth rates, LDM schedules result in increased tumor cell reduction.*

Since most tumors do not grow at a fixed exponential rate throughout their entire growth history [5, 6], a more generally applicable model, developed subsequently by Norton and Simon [7], is based on Gompertzian growth of a tumor [6,8], which is exponential growth, but with an exponentially decaying growth rate 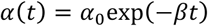 The decaying growth rate can be viewed partly as a simple consequence of the fact that the ratio of tumor surface area (SA) to volume (V) scales like 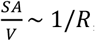, for a spherical tumor of radius R. Since tumors need to supply their volume with nutrients and oxygen which they (mostly) accomplish via diffusion through their outer surface, as R increases, this becomes more challenging causing the tumor growth to slow down (although there are certainly strategies the tumors develop to partly mitigate this effect). Smaller regression rates for these larger, more slow growing tumors is also consistent with this scaling law since lethal toxins enter the tumor core primarily through its surface. For *β* = 0 the Gompertzian model reduces to the previous model of Skipper and Schabel. The new model gives rise to Gompertzian sigmoidal shaped growth curve [9] shown in Figure 1 for three different tumor types (slow growth, medium growth, fast growth). The Norton-Simon hypothesis (now viewed as a bedrock principle) says that tumor cell reduction, if administered starting at time *t*_1_, is proportionalto the *unperturbed* growth rate α(*t*_1_)Thus it references the *instantaneous* growth rate of the *unperturbed* tumor at the time one cycle of chemotherapy is initiated [7,10,11].

Examples of tumor cell reduction are shown in Figure 1. Since the timescale on which a single cycle of chemotherapy is administered is short compared to the timescale on which a tumor size changes significantly, the (quasi-steady) approximation is not unreasonable and can be viewed as a generalization of Skipper’s laws for time-dependent growth rates. However with the potential application of several cycles starting at times *t*_2_,*t*_3_ … perhaps of varying lengths and doses (metronomic vs. maximum tolerated dose [12,13]), and with unperturbed growth rates and tumor sizes differing at the beginning and end of each cycle, the quantitative efficacy of this approximation begins to break down. The question then arises as to what would be an optimal chemotherapeutic scheduling strategy (varying concentration, dose, density) that accommodates the full complex dynamical range of features associated with tumor cell kinetics?

## II. ILLUSTRATIVE RESULTS OF APPLICATION OF METHODS

The total dose, D, of any arbitrary chemotherapy schedule is the sum of the nondimensional doses, *c_i_* (*0* ≤ *c* ≤ *1*), on each of the *i* days (some of which can be zero). *N* is the number of days in a given cycle (also known as the intercycle time):

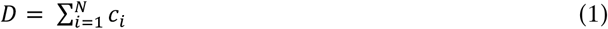

The dose density, d, of a regimen can be found by summing the number of days where a non-zero dose is delivered, and dividing by the intercycle time in days, *N*. The density is a non-dimensional parameter such that *0* ≤ *d* ≤ *1*.

The quasi-steady approximation of the Norton-Simon hypothesis can be extended to find optimal chemotherapy schedules over the full course of treatment (see [14]). Norton and Simon relate tumor regression to the total dose delivered, but the relationship between dose (ci) and density (d) is not made clear by the hypothesis. The relationship between dose and density has been studied by Prokopiou [15] (termed “fractionation”) in radiotherapy by using the proliferation saturation index (PSI, the ratio of volume to tumor carrying capacity) to predict radiation response and to personalize optimal radiotherapy schedules. The PSI (inversely related to the growth rate) of the tumor is found to be a good prognostic factor for radiation response. Figure 2 depicts a series of chemotherapeutic strategies quantified by the Shannon entropy, E, of the strategy (2), for a range of dose, density, and entropy values [16]. Low-dose metronomic strategies generally correspond to high-entropy strategies, whereas maximum tolerated dose strategies correspond to low-entropy strategies. In Figure 3 we show a histogram of all possible different entropy-based strategies, plotted as a function of tumor-cell reduction. The graph shows a clear tendency for higher entropy strategies to produce greater tumor-cell reduction. This is discussed in more detail in [17]. Entropy is calculated from the formula:

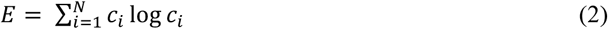

Figure 4 points to the fact that over multiple cycles, higher entropy strategies have a bigger impact on faster growing tumors (purple) than on slower growing tumors (blue), as evidenced by the larger slope of the regression line.

**Figure 2:**
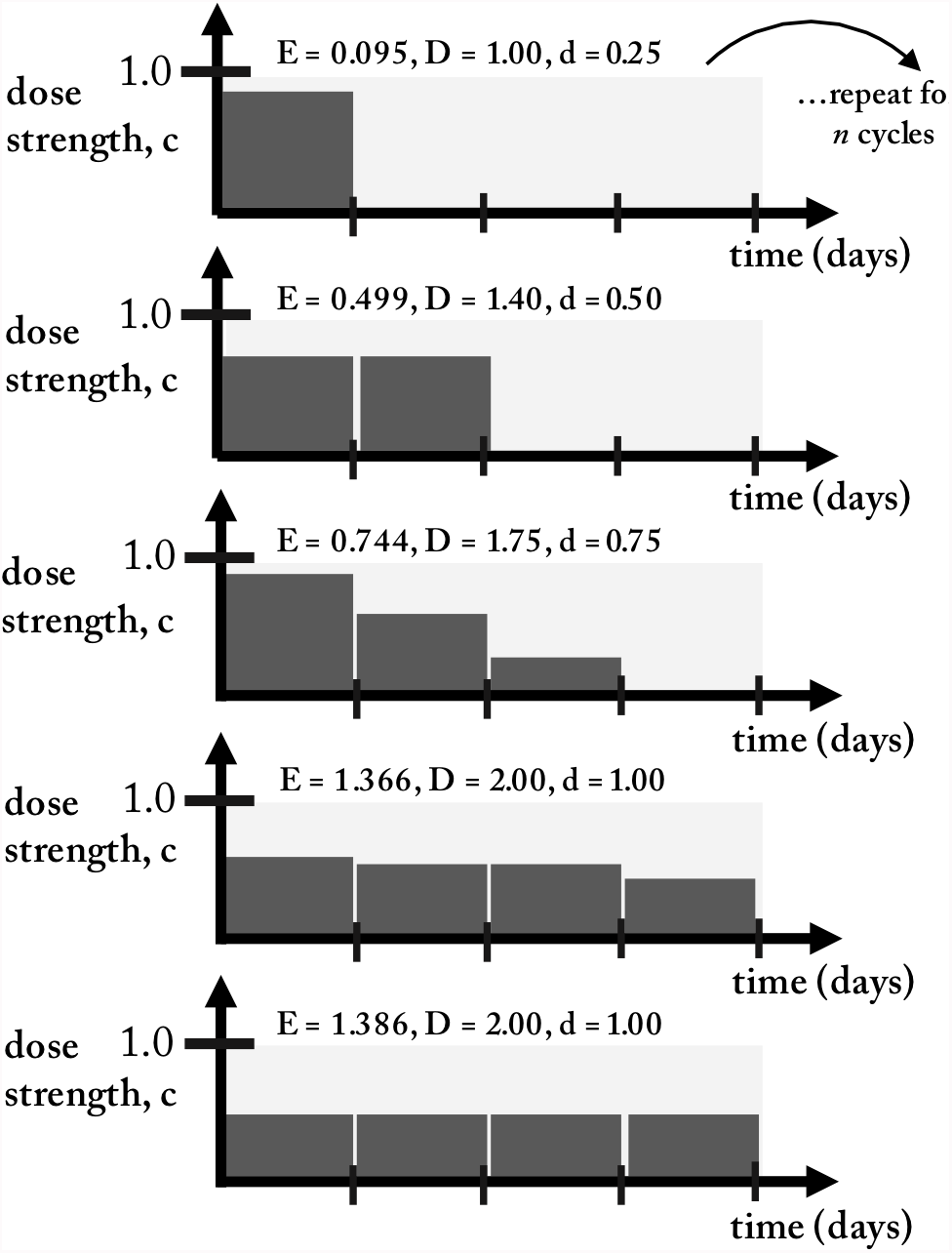
Shannon entropy as an index to compare sample 1-cycle 4-day treatment strategies. *Chemotherapy regimens can be simulated for a range of dose, density, and entropy values. Pictured from top to bottom are a range of representative regimens from low entropy (i.e. high dose, low density) to high entropy (i.e. low dose, high density) for a cycle of N = 4 days. On each i^th^ day, treatment of dose *c*_*i*_ is administered. The treatment strategy’s Shannon entropy, E, is calculated according to equation 2 and the total dose, D, is the sum of all daily doses (equation 1). It should be noted that LDM-like regimens correspond to a high entropy value (e.g. botom graph).*

**Figure 3:**
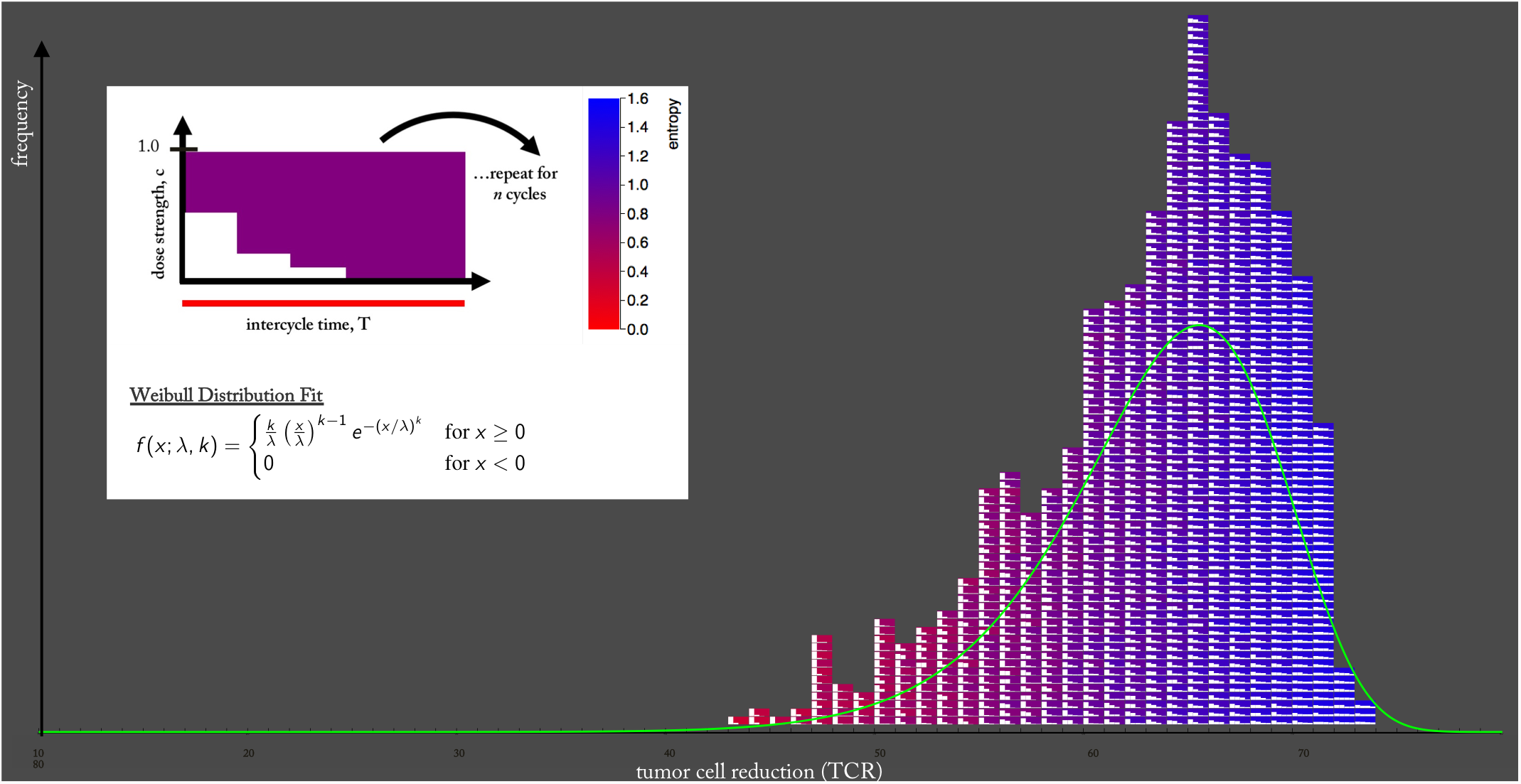
High entropy, LDM-like chemotherapies outperform low entropy MTD-like chemotherapies. *A pictorial histogram is plotted where each chemotherapy schedule pictorial representation (colorcoded from red: low entropy to blue: high entropy) is arranged by tumor cell reduction (TCR). All regimens are equivalent total dose (D = 0.3), monotonically decreasing, and are repeated for 8 cycles of chemotherapy for a slow growing tumor (w = 0.1). The histogram clearly shows a color-shift from red toward blue for low TCR, ineffective therapies toward high TCR, effective therapies. High entropy therapies outperform low entropy therapies. The data was fit to a Weibull distribution (equation shown in panel inset; k = 14.251, λ = 65.882), overlaid in green.*

**Figure 4:**
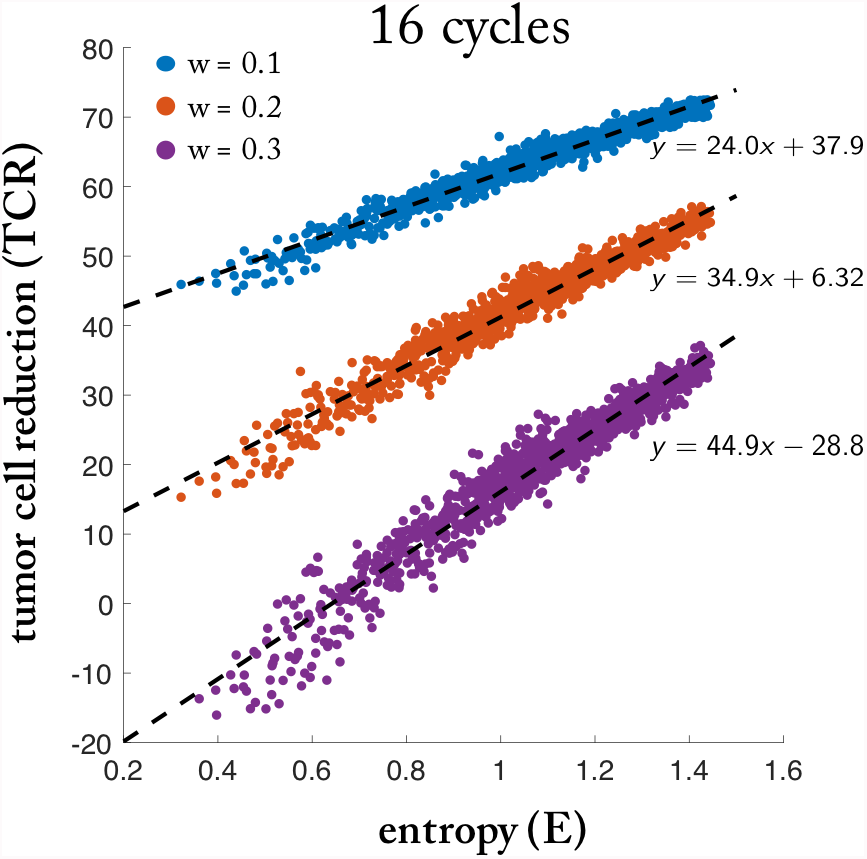
High entropy strategies lead to an increase in tumor regression. TCR, (averages of 25 stochastic simulations for total dose delivered D = 0.3) and entropy (E) is shown for a 16 cycles of chemotherapy, repeated for slow (w = 0.1, blue), medium (w = 0.2, red), and fast growing tumors (w = 0.3, purple). High entropy strategies outperform low entropy. Fast growing tumors (purple) has the highest slope (dashed lines) highlighting the relative advantage of high entropy strategies for higher tumor growth rates.

The concept of choosing dosing schedules and strategies based on tumor growth rates is not currently done in medical practice and might prove to be a fruitful idea to test further in clinical trials focused on this question. Mathematical models and computer simulations can play an important supportive role in helping to predict possible responses and in forming hypotheses to test further.

## III. QUICK GUIDE TO THE METHODS

The mathematical model used for the results shown is based on a finite-cell stochastic Moran process [18,19] with cell-cell interactions based on a Prisoner’s Dilemma payoff matrix A:

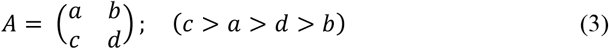

Fitness of the healthy cells H (cooperators), and the cancer cells C (defectors) are given by:

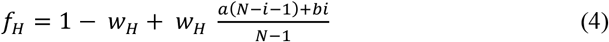

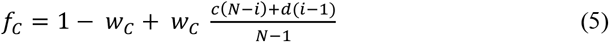

The parameters *w*_*H*_ and *w*_*C*_ (*0* ≤ *w* ≤ *1*) measure the strength of the selection pressure on each population of cells, Selection is initially equivalent (*w*_*H*_ = *w*_*C*_ = w), and altered by the dose concentration, *c*, during therapy as follows:

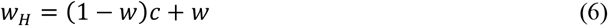

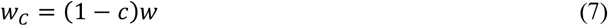

If there are ‘*i*’ cancer cells and *N-i* healthy cells (*N* is the total population of cells), then the transition probabilities at each step in the Moran process are given by [20]:

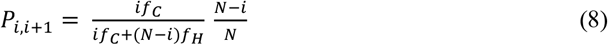

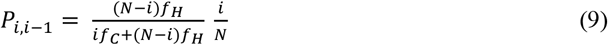

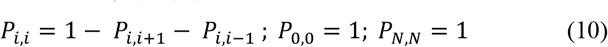

